# A new machine learning based computational framework identifies therapeutic targets and unveils influential genes in pancreatic islet cells

**DOI:** 10.1101/2022.05.01.490186

**Authors:** Turki Turki, Y-h. Taguchi

## Abstract

Pancreatic islets comprise a group of cells that produce hormones regulating blood glucose levels. Particularly, the alpha and beta islet cells produce glucagon and insulin to stabilize blood glucose. When beta islet cells are dysfunctional, insulin is not secreted, inducing a glucose metabolic disorder. Identifying effective therapeutic targets against the disease is a complicated task and is not yet conclusive. To close the wide gap between understanding the molecular mechanism of pancreatic islet cells and providing effective therapeutic targets, we present a computational framework to identify potential therapeutic targets against pancreatic disorders. First, we downloaded three transcriptome expression profiling datasets pertaining to pancreatic islet cells (GSE87375, GSE79457, GSE110154) from the Gene Expression Omnibus database. For each dataset, we extracted expression profiles for two cell types. We then provided these expression profiles along with the cell types to our proposed constrained optimization problem of a support vector machine and to other existing methods, selecting important genes from the expression profiles. Finally, we performed (1) an evaluation from a classification perspective which showed the superiority of our methods against the baseline; and (2) an enrichment analysis which indicated that our methods achieved better outcomes. Results for the three datasets included 44 unique genes and 10 unique transcription factors (SP1, HDAC1, EGR1, E2F1, AR, STAT6, RELA, SP3, NFKB1, and ESR1) which are reportedly related to pancreatic islet functions, diseases, and therapeutic targets.

## 1. Introduction

Pancreatic islets comprise several endocrine cells that play a key role in metabolism [1, 2]. Among these, alpha and beta islet cells are endocrine cells involved in regulating blood glucose levels [3]. As carbohydrates break down into glucose and enter the blood stream, beta cells secrete insulin to absorb glucose into cells for energy. Insulin mainly aims to store excess glucose in the liver [4–6]. When beta cells do not function properly, insulin is not secreted and blood glucose levels are elevated. Left untreated, this condition can lead to the development of diabetic complications, such as blindness, cancer, and kidney damage, among others [7, 8]. Advances in biology and medical research have led to an improved understanding of the pancreatic functions, diseases, and therapeutic targets.

Zheng et al. [9] downloaded transcriptome expression profiling dataset (GSE119794) pertaining to pancreatic adenocarcinoma (PAAD) from the GEO database and used a bioinformatics approach to identify 262 differentially expressed genes between patients and control profiles through a limma package in R. Then, an enrichment analysis was applied in which three hub genes (CXCL5, KRT19, and COL11A1) were identified by cytoHubba from the protein-protein interactions (PPI) network. The three hub genes were submitted to GEPIA and TIMER within the TCGA database. Results indicated that COL11A1 was highly correlated and expressed in many expression profiles pertaining to PAAD, suggesting that the gene could act as a biomarker in PAAD.

Prashanth et al. [10] proposed a bioinformatics approach to identify hub genes that may play a key role in understanding type 1 diabetes (T1D) and act as targets in the treatment and diagnosis of the disease. First, they downloaded an expression-profiling dataset (GSE123658) consisting of 39 and 43 expression profiles for T1D and healthy individuals, respectively, from the GEO database. Totally, 284 differentially expressed genes were identified through the limma package in R, out of which an enrichment analysis identified 10 hub genes (EGFR, POLR2A, PRKACA, GRIN2B, GABARAP, TLN1, GJA1, CAP2, MIF, and PXN) [10]. Kang et al. [11] aimed to identify genes that can act as a therapeutic targets of T2D patients with periodontitis comorbidity. Five expression profiling datasets were downloaded from the GEO database and a total of 152 commonly DEGs were identified between T2D and periodontitis patients. An enrichment analysis further identified four hub genes (PTPRC, HGF, RAC2, and INPP5D) from the PPI network, which can act as therapeutic targets for T2D patients with periodontitis. Chan et al. identified common driver genes for T2D and cardiovascular disease from genome-wide association studies within women of different ethnicities [12]. Other studies have also contributed to the understanding of molecular mechanism of pancreas-related disorders and provide therapeutic targets [13–17].

Several computational methods have been developed for the identification of significant differentially expressed genes as inputs for enrichment analysis tools (e.g., Enrichr) to evaluate biological functions. For example, Taguchi et al. [1] proposed an unsupervised feature extraction method to identify drug candidates for COVID-19. The proposed method, which utilizes data represented as 4D tensors, was applied to gene expression profiles for SARS-CoV-2-infected lung cancer cell lines, generating 163 genes related to the SARS-CoV-2 infection process. Enrichr was then used to identify drug compounds expected to alter the expression of the selected genes. The proposed approach outperformed other bioinformatics-based approaches. Taguchi et al. [2] integrated several gene expression profiles to represent data as 6D tensors for unsupervised feature extraction based on tensor decomposition. An Enrichr analysis of the 2281 genes identified by the method successfully revealed various diseases associated with diabetes. Other unsupervised feature extraction methods aimed at extracting biological information about selected genes have also been proposed [18, 19].

Although current advances in biology and medical research have contributed to a better understanding of pancreatic disorders and diseases, advanced computational frameworks are needed to uncover the genetic architecture of various metabolic disorders. These frameworks will enable us to identify influential genes that play key roles in pancreatic islet function and identify therapeutic targets. They will advance our understanding of shared molecular mechanisms. The process of drug discovery can be accelerated by computational frameworks because they reduce the search space of potential therapeutic targets.

### Contributions

We list our main contributions as follows:

1. We present a computational framework comprising a novel machine learning formulation of support vector machines (SVM) [20], followed by biological enrichment analysis, to retrieve influential genes that will allow us to (a) understand the molecular mechanisms of the pancreas; (b) improve targeted therapy related to the pancreas; and (c) discover common molecular mechanisms in different cell types.
2. We compare the proposed framework with other popular methods, including SVM, limma [21], significance analysis of microarrays (SAM), and *t*-test [22] for the identification of genes related to pancreatic islet functions, diseases and therapeutic targets.
3. We evaluate all methods from two perspectives, using three transcription profiling datasets pertaining to pancreatic cells (mainly islet cells) from the GEO database. From the biological perspective, experiments identified 44 genes and 10 transcription factors; these have been reported to be related to pancreatic functions, diseases, and therapeutic targets. From the classification perspective, we demonstrate that our new formulation of SVM contributes to improved performance results when incorporated into our computational framework.

### Organization

The rest of the study is organized as follows. In Section 2, we describe the transcriptome expression profiling datasets used in this study, followed by our proposed computational framework. In Section 3, we evaluate and report the performance results of all methods considered in this study and discuss these in Section 4. Finally, we provide a conclusion to this work, pointing out future directions.

## 2. Materials and Methods

### 2.1 Transcriptome Expression Profiles

We downloaded three single-cell RNA-seq datasets from the NCBI GEO according to the following GEO IDs.

#### 2.1.1 GSE87375: Dataset1

We retrieved this single-cell dataset using the GEO ID GSE87375 [23], considering 913 single-cells and 40916 expression values. Therefore, we have a 913 x 40917 matrix including a column vector for the cell type. The 913 cells comprise 338 alpha and 575 beta cells in the pancreatic islets. We refer to the remaining dataset as Dataset1.

#### 2.1.2 GSE79457: Dataset2

This dataset comprised 181 single-cells and 23431 expression values [24] encoded as a 181 x 23432 matrix including the cell type column vector. The single-cells comprised 63 alpha and 31 delta cells, 47 beta cells and 40 other cells. We refer to this dataset as dataset2. It is noteworthy that although the original dataset had 182 single-cells, one expression profile was excluded because it was defective.

#### 2.1.3 GSE110154: Dataset3

The third dataset comprised 1860 single-cells and 22762 expression values [25]; 376 cells were acinar and 42 were delta pancreatic cells. The 1442 remaining cells were other types. We refer to this dataset in the remaining sections as dataset3.

### 2.2 Computational Framework

In Figure 1, we show the steps carried out by the proposed computational framework. Suppose we have a single-cell dataset as {(x_1_,y_1_),…,(x_m_,y_m)_}, where x_i_ represents the i*th* cell and y_i_ is the corresponding cell type, in this study, we consider y_i_ in the binary case. We encode all cells x_*i*_ (for *i* = 1..m) in this dataset as an *m* × *n* matrix, where *m* and *n* are the number of cells and genes, respectively. Similarly, we encode all cell types y_*i*_ (for *i* = 1..m) as a 1 × *m* column vector. We seek to identify *p* influential genes out of the *n* genes, such that these *p* genes are mainly expressed in a given cell type. As a result, we propose a robust variant of the SVM based on the HR (See Supplementary Additional File 1: Equations in S1.1 and S1.2), working as follows. In Equation 1, the constrained optimization problem finds the weights w = [*w*_1_…*w*_n_] (a vector consisting of *n* elements) and bias *b* ∊ R (consisting of one value) such that the cost function is minimized. As each cell is encoded by *n* genes, the weights w correspond to the importance of these genes among all cells. The higher the weight *w*_i_ is, the more important the gene *i* is.

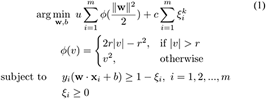

**Figure 1:**
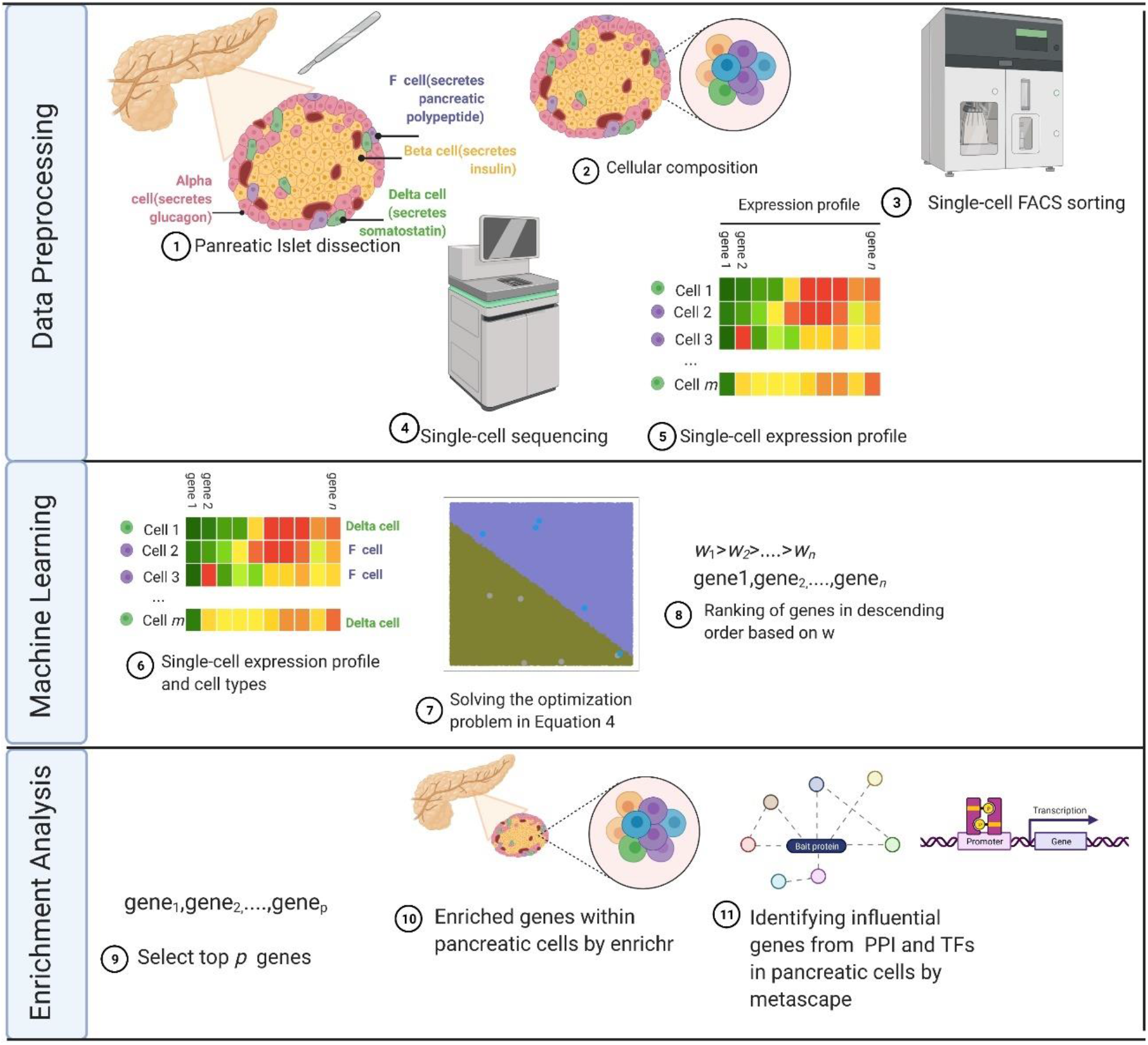
Flowchart of the proposed computational framework to identify influential genes expressed in pancreatic islet cells. Figure created with BioRender.com.

According to Equation 1, the constrained optimization problem becomes as shown in Equation 2 when |*v*| > *r*. Otherwise, the optimization problem becomes as in Equation 3. Depending on the dataset, this leads to different cost functions and different weights as a result, when compared to the original SVM in Supplementary Additional File 1. To solve both constrained optimization problems in Equations 2 and 3 and obtain weights w and bias b, we utilize the CVXR solver in R [26].

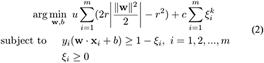

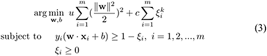

## 3 Experiments and Results

### 3.1 Experimental Methodology

We compare our proposed methods (i.e., scsvm1 and scsvm2) against the baseline and other well-known computational methods, such as limma [21], SAM, and *t*-test. [22]. All the approaches work under the supervised learning approach, where each method takes labeled examples to induce the model. For methods using SVM (i.e., scsvm1, scsvm2 and baseline), because the SVM model is expressed as weighted features plus the bias (i.e., w.x + *b*), we select the top *p* genes based on the top *p* values in w, as those correspond to genes with high importance. For the remaining methods, we select significant genes according to adjusted *p*-values < 0.01. In Table 1, we give a summary of the methods used in this study.

**Table 1:** Summary of studied Computational Methods.

| Abbreviation | Prediction Algorithm |
| --- | --- |
| scsvm1 | The first single-cell SVM |
| scsvm2 | The second single-cell SVM |
| baseline | The baseline SVM (See Supplementary Additional File 1: Section S1.1) |
| limma | Linear models for microarray data |
| sam | Significance analysis of microarrays |
| *t*-test | Student's *t*-test |
| lasso | Least absolute shrinkage and selection operator |

To evaluate the performance results, we performed a biological enrichment analysis by uploading genes of each method to Enrichr (https://maayanlab.cloud/Enrichr/) [27] and Metascape (https://metascape.org/gp/index.html) [28]. Assuming the term is a cell, the greater the expressed genes in a cell, the better the performance when the cell is related to the studied cell type. The same holds when a term is a tissue (i.e., consisting of cells performing the same function). Moreover, the lower the ranking of the term (e.g., cell), the better the performance. We also evaluate the models from a classification perspective, using area under curve (AUC) performance measures, where the higher the AUC, the better the performance. We performed experiments in this study using R [29]. Specifically, we used the CVXR package in R to solve optimization problems [26]. The siggenes package in R was used for performing SAM [30]. The lmFit and eBayes functions in the limma package in R were utilized for model fitting [21] and the t.test function in the R stats package was used for comparisons [29]. The glmnet package in R was used for performing lasso [31] and selecting genes associated with non-zero coefficients as these denote important genes. All four methods (i.e., SAM, limma, *t*-test, and lasso) utilize expression profiles and corresponding cell types. For the three methods (i.e., SAM, limma, and *t*-test), the p.adjust function with the “BH” option was utilized to compute adjusted *p*-values, setting a threshold of *p* < 0.01, as in [32].

### 3.2 Results

#### 3.2.1 Dataset1

Table 2 shows terms identified using Enrichr. Related terms were ranked highly and associated with a large number of expressed genes, demonstrating the validity of the computational method. It can be shown from Table 2 that the top three retrieved terms (when scsvm1 is utilized) are pancreatic islets, beta cells, and alpha cells (written in bold in the overlap column). Thirty-nine out of 95 genes were expressed in the pancreatic islets and distributed into its cells. Twenty-nine of the 95 enriched genes were expressed in pancreatic beta and alpha cells. Twenty-seven genes (PCSK2, SLC7A14, SLC7A2, TMEM27, PCSK1N, TTR, EPCAM, PEG10, NPY, TSPAN7, PPY, CHGB, CHGA, PEG3, GPX3, ETV1, KCNK16, PAX6, TM4SF4, SMARCA1, ISL1, PYY, G6PC2, SCG5, CPE, IAPP, GHRL) out of these 29 were common between alpha and beta islet cells. The *p-*values indicate that these outcomes are significant. The second-best method is scsvm2, where the top two retrieved items are pertaining to pancreatic islets and beta cells. The term alpha cell ranked sixth. These results demonstrate the superiority of scsvm1 and scsvm2, when compared to all other methods where all cells pertaining to the pancreas are not in the top three items with more expressed genes. Although the top retrieved terms in lasso are pancreatic islets, beta cells, and alpha cells, fewer genes were expressed. Regarding Dataset 1, we provide a list of genes obtained via each method in Supplementary Data Sheet 1. Moreover, we provide a full list of enrichment analysis results of each method in Supplementary Table 1.

**Table 2:** Enriched terms in “ARCHS4 Tissues” using Enrichr based on genes obtained from Dataset1, consisting of alpha and beta pancreatic cells. Rank column shows the order of terms when retrieved. Overlap displays the number of common genes between the uploaded gene and those in the term divided by the number of genes in that term. Best results of a method is in bold.

| Method | Rank | Term | Overlap | P-value | Adjusted P-value |
| --- | --- | --- | --- | --- | --- |
| scsvm1 | 1 | PANCREATIC ISLET | **39/2316** | $2.23 \times 10^{-13}$ | $2.41 \times 10^{-11}$ |
| | 3 | BETA CELL | **29/2316** | $5.79 \times 10^{-7}$ | $2.08 \times 10^{-5}$ |
| | 2 | ALPHA CELL | **29/2316** | $5.79 \times 10^{-7}$ | $2.08 \times 10^{-5}$ |
| scsvm2 | 1 | PANCREATIC ISLET | 34/2316 | $6.31 \times 10^{-10}$ | $6.56 \times 10^{-8}$ |
| | 2 | BETA CELL | 24/2316 | $1.60 \times 10^{-4}$ | $8.30 \times 10^{-3}$ |
| | 6 | ALPHA CELL | 20/2316 | $5.60 \times 10^{-3}$ | $8.33 \times 10^{-2}$ |
| baseline | 1 | PANCREATIC ISLET | 33/2316 | $2.71 \times 10^{-9}$ | $2.81 \times 10^{-7}$ |
| | 5 | BETA CELL | 22/2316 | $1.06 \times 10^{-3}$ | $1.83 \times 10^{-2}$ |
| | 8 | ALPHA CELL | 19/2316 | $1.19 \times 10^{-2}$ | $1.23 \times 10^{-1}$ |
| limma | 101 | PANCREATIC ISLET | 4/2316 | $9.97 \times 10^{-1}$ | $1.00 \times 10^0$ |
| | 46 | BETA CELL | 11/2316 | $5.48 \times 10^{-1}$ | $1.00 \times 10^0$ |
| | 83 | ALPHA CELL | 7/2316 | $9.34 \times 10^{-1}$ | $1.00 \times 10^0$ |
| sam | 3 | PANCREATIC ISLET | 16/2316 | $7.91 \times 10^{-2}$ | $1.00 \times 10^0$ |
| | 1 | BETA CELL | 16/2316 | $7.91 \times 10^{-2}$ | $1.00 \times 10^0$ |
| | 10 | ALPHA CELL | 11/2316 | $5.48 \times 10^{-1}$ | $1.00 \times 10^0$ |
| t-test | 50 | PANCREATIC ISLET | 8/2316 | $8.73 \times 10^{-1}$ | $1.00 \times 10^0$ |
| | 2 | BETA CELL | 14/2316 | $2.07 \times 10^{-1}$ | $1.00 \times 10^0$ |
| | 28 | ALPHA CELL | 9/2316 | $7.85 \times 10^{-1}$ | $1.00 \times 10^0$ |
| lasso | 3 | PANCREATIC ISLET | 4/2316 | $8.53 \times 10^{-3}$ | $1.39 \times 10^{-1}$ |
| | 1 | BETA CELL | 5/2316 | $8.58 \times 10^{-4}$ | $4.20 \times 10^{-2}$ |
| | 2 | ALPHA CELL | 4/2316 | $8.53 \times 10^{-3}$ | $1.39 \times 10^{-1}$ |

Because scsvm1 exhibited good results in terms of retrieving the cells related to studying trancriptome expression profiles in dataset1, we uploaded 95 genes of scsvm1 to Metascape. Figure S2.1 in Supplementary Additional File 2 shows the protein-protein interaction (PPI) network obtained from Metascape. Figure 2 shows the highly connected subgraphs obtained from the PPI network, which include the following 16 genes: PPY, PYY, NPY, GNG12, BSG, HSP90AA1, HSPA8, RBM39, H1-2, HSP90AB1, RRM2, TUBBB, ATP5F1B, ACTB, SLC25A5, and EEF1A1.

**Figure 2:**
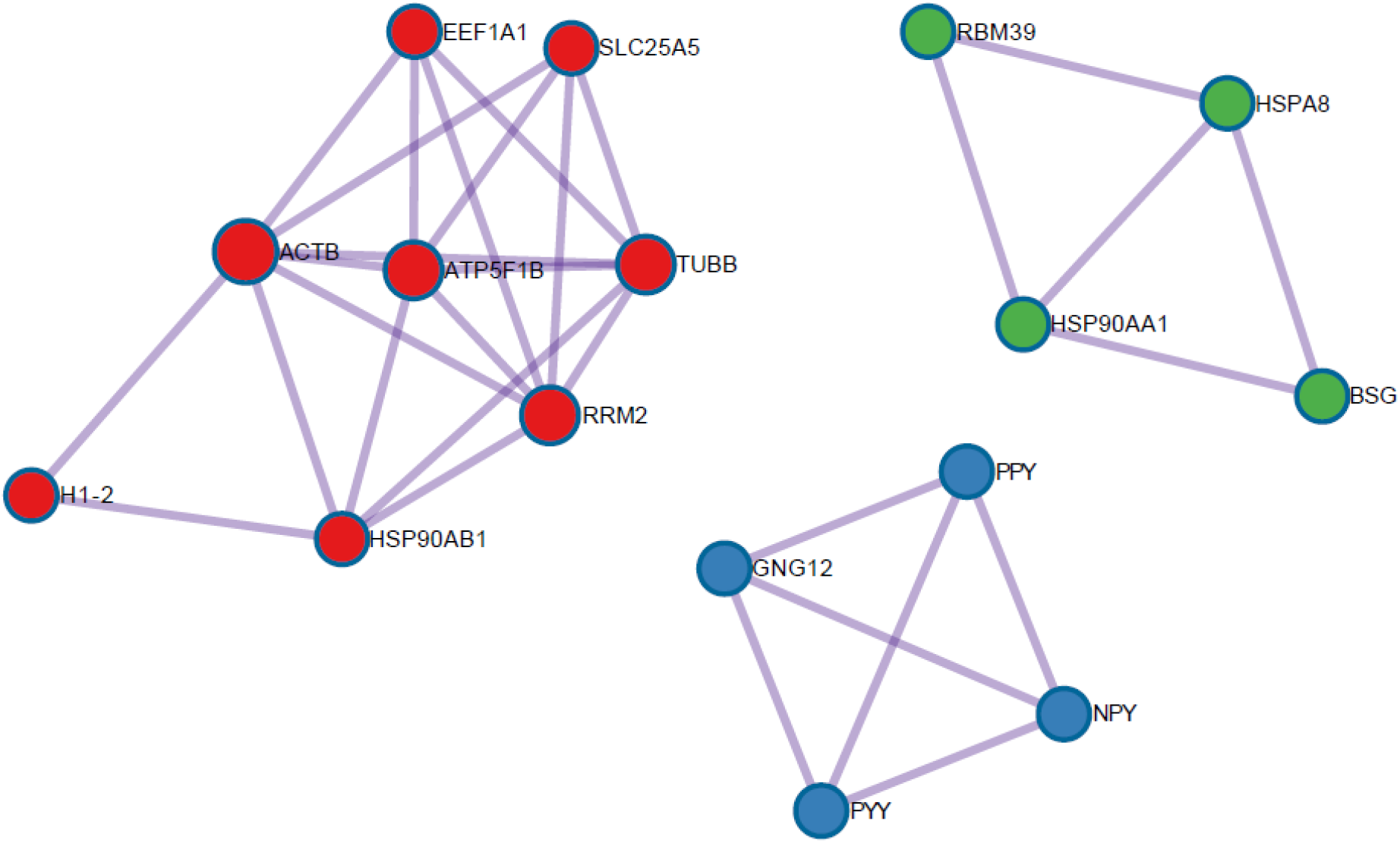
Three highly interconnected regions (clusters) in a protein-protein interaction enrichment analysis according to 95 genes of scsvm1, submitted to metascape.

Some of these identified genes have played a key role in understanding the working mechanism of pancreatic cells [33, 34]. In Figure S2.2 in Supplementary Additional File 2, we identified four transcription factors-SP1, HDAC1, EGR1 and E2F1. SP1 has been reported as a biomarker for identifying the most common type of pancreatic cancer-pancreatic ductal adenocarcinoma [35]. HDAC1 has also been reported to control the expression of the p53 gene in pancreatic cancer. Additionally, EGR1 and E2F1 control the expression of cancer cells [36, 37]. These 16 identified genes and their 4 transcription factors can aid as therapeutic targets pertaining to common disorders in the pancreas.

#### 3.2.2 Dataset2

Table 3 reports the retrieved terms for the genes uploaded by each method to Enrichr. It can be shown that our method scsvm2 outperforms all other methods. Specifically, pancreatic islet is ranked at the top, with 20 expressed genes out of 2316. Thirteen and twelve genes were expressed in beta and alpha cells, respectively. Eleven genes (PCSK2, MPP2, PEG3, ABCC8, GCG, BAIAP3, EML5, NR4A1, ARC, GNG4, and CHGB) were commonly expressed in beta and alpha cells. In terms of the number of genes, sam, *t*-test, and lasso differed from scsvm2 and scsvm1. The results reflect that scsvm2 had more expressed genes in these terms than all other methods (see results in bold in the overlap column). Results obtained by scsvm1 and baseline are almost the same. Although SAM generated the highest-ranking results and a number of significantly expressed genes, fewer expressed genes were obtained by this approach than by scsvm1 and scsvm2. This can be explained by the dependence of significance scores on the input gene list based computational method and the gene list for the retrieved terms. Therefore, all genes obtained by SAM overlap with the genes in the term list.

**Table 3:**
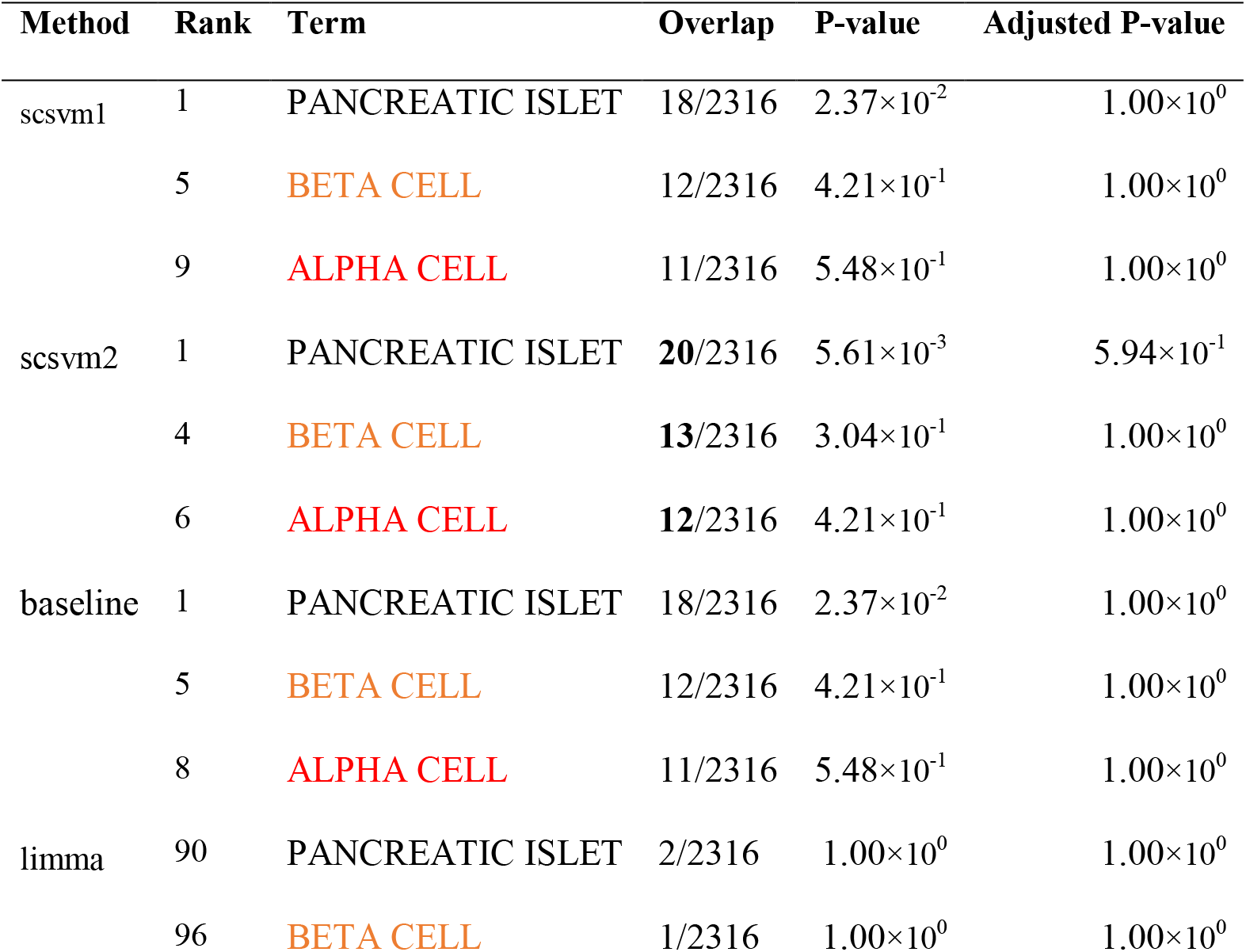
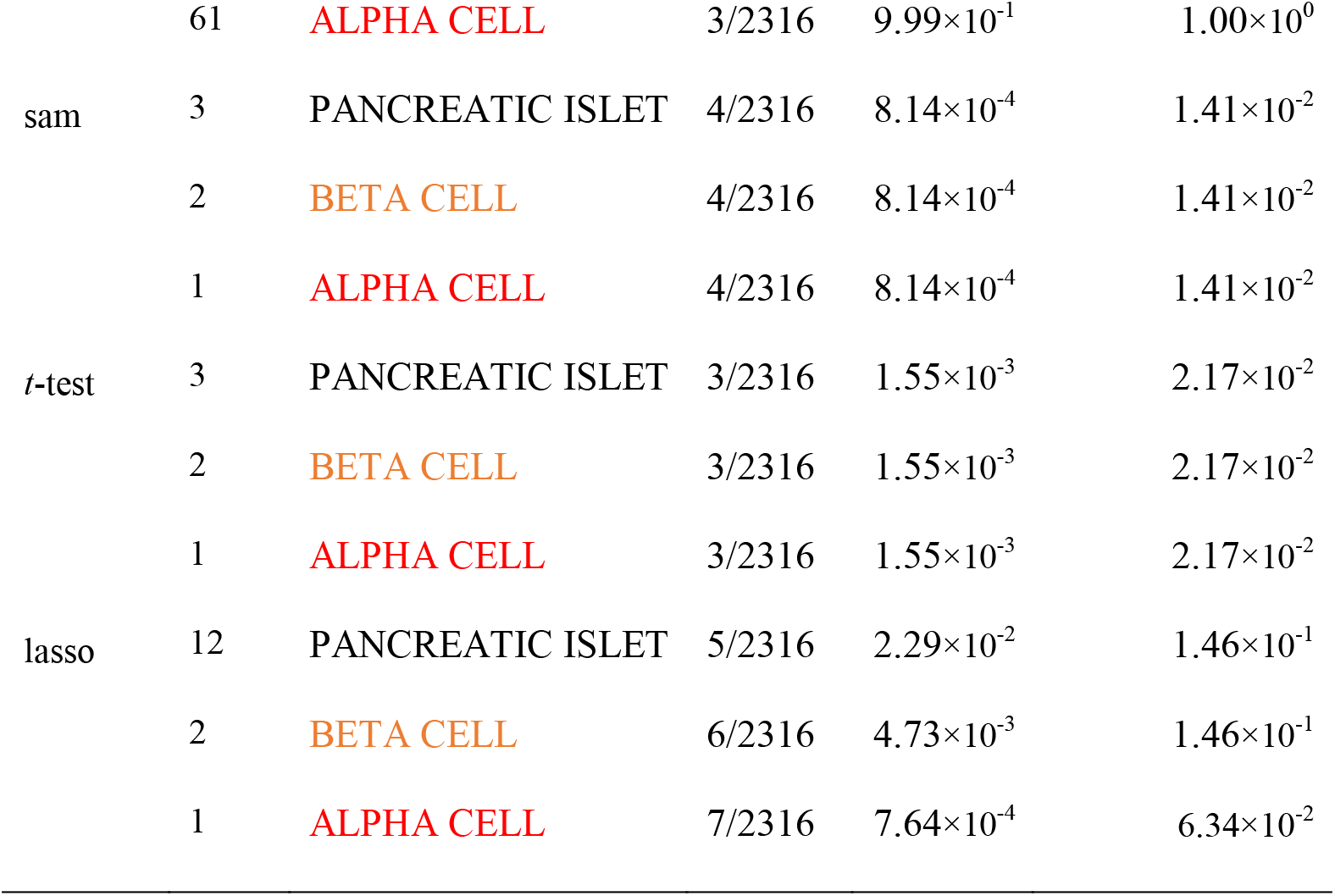
Enriched terms in “ARCHS4 Tissues” using Enrichr based on genes obtained from Dataset2, consisting of alpha and delta pancreatic cells. Rank column shows the order of terms when retrieved. Overlap displays the number of common genes between the uploaded gene and those in the term divided by the number of genes in that term. Best results of a method is in bold.

| Method | Rank | Term | Overlap | P-value | Adjusted P-value |
| --- | --- | --- | --- | --- | --- |
| scsvm1 | 1 | PANCREATIC ISLET | 18/2316 | $2.37 \times 10^{-2}$ | $1.00 \times 10^0$ |
| | 5 | BETA CELL | 12/2316 | $4.21 \times 10^{-1}$ | $1.00 \times 10^0$ |
| | 9 | ALPHA CELL | 11/2316 | $5.48 \times 10^{-1}$ | $1.00 \times 10^0$ |
| scsvm2 | 1 | PANCREATIC ISLET | **20**/2316 | $5.61 \times 10^{-3}$ | $5.94 \times 10^{-1}$ |
| | 4 | BETA CELL | **13**/2316 | $3.04 \times 10^{-1}$ | $1.00 \times 10^0$ |
| | 6 | ALPHA CELL | **12**/2316 | $4.21 \times 10^{-1}$ | $1.00 \times 10^0$ |
| baseline | 1 | PANCREATIC ISLET | 18/2316 | $2.37 \times 10^{-2}$ | $1.00 \times 10^0$ |
| | 5 | BETA CELL | 12/2316 | $4.21 \times 10^{-1}$ | $1.00 \times 10^0$ |
| | 8 | ALPHA CELL | 11/2316 | $5.48 \times 10^{-1}$ | $1.00 \times 10^0$ |
| limma | 90 | PANCREATIC ISLET | 2/2316 | $1.00 \times 10^0$ | $1.00 \times 10^0$ |
| | 96 | BETA CELL | 1/2316 | $1.00 \times 10^0$ | $1.00 \times 10^0$ |
| | 61 | ALPHA CELL | 3/2316 | $9.99 \times 10^{-1}$ | $1.00 \times 10^0$ |
| sam | 3 | PANCREATIC ISLET | 4/2316 | $8.14 \times 10^{-4}$ | $1.41 \times 10^{-2}$ |
| | 2 | BETA CELL | 4/2316 | $8.14 \times 10^{-4}$ | $1.41 \times 10^{-2}$ |
| | 1 | ALPHA CELL | 4/2316 | $8.14 \times 10^{-4}$ | $1.41 \times 10^{-2}$ |
| t-test | 3 | PANCREATIC ISLET | 3/2316 | $1.55 \times 10^{-3}$ | $2.17 \times 10^{-2}$ |
| | 2 | BETA CELL | 3/2316 | $1.55 \times 10^{-3}$ | $2.17 \times 10^{-2}$ |
| | 1 | ALPHA CELL | 3/2316 | $1.55 \times 10^{-3}$ | $2.17 \times 10^{-2}$ |
| lasso | 12 | PANCREATIC ISLET | 5/2316 | $2.29 \times 10^{-2}$ | $1.46 \times 10^{-1}$ |
| | 2 | BETA CELL | 6/2316 | $4.73 \times 10^{-3}$ | $1.46 \times 10^{-1}$ |
| | 1 | ALPHA CELL | 7/2316 | $7.64 \times 10^{-4}$ | $6.34 \times 10^{-2}$ |

Because scsvm2 had more expressed genes in pancreatic cells, we uploaded these 95 genes by scsvm2 to Metascape. In Figure S2.3 in Supplementary Additional File 2, we show the PPI network provided by Metascape. Two highly connected subgraphs are obtained as shown in Figure 3, which include the following 14 genes: ITG1, HSPA5, ACTB, ATP5F1B, CCT5, GAPDH, ARPC1A, GNA12, ARFGAP1, INS, CDC42, BSG, ARFGAP3, and ARF1. Among these 14 genes, it has recently been reported that ARFGAP1 is a predictive factor for pancreatic cancer [38], and GAPDH has been used as a reference gene in fetal rat pancreas analysis [39]. In Figure S2.4 in Supplementary Additional File 2, only the androgen receptor (AR) transcription factor was reported by Metascape. It has been reported recently that AR plays a key role in the progression of pancreatic cancer [40]. These results demonstrate that the inhibition of AR and the understanding of these 14 genes can act as a therapeutic target in pancreatic cancer. In terms of dataset 2, a list of genes obtained via each method is available in Supplementary Data Sheet 2. Additionally, a full list of enrichment analysis results of each method is available in Supplementary Table 2.

**Figure 3:**
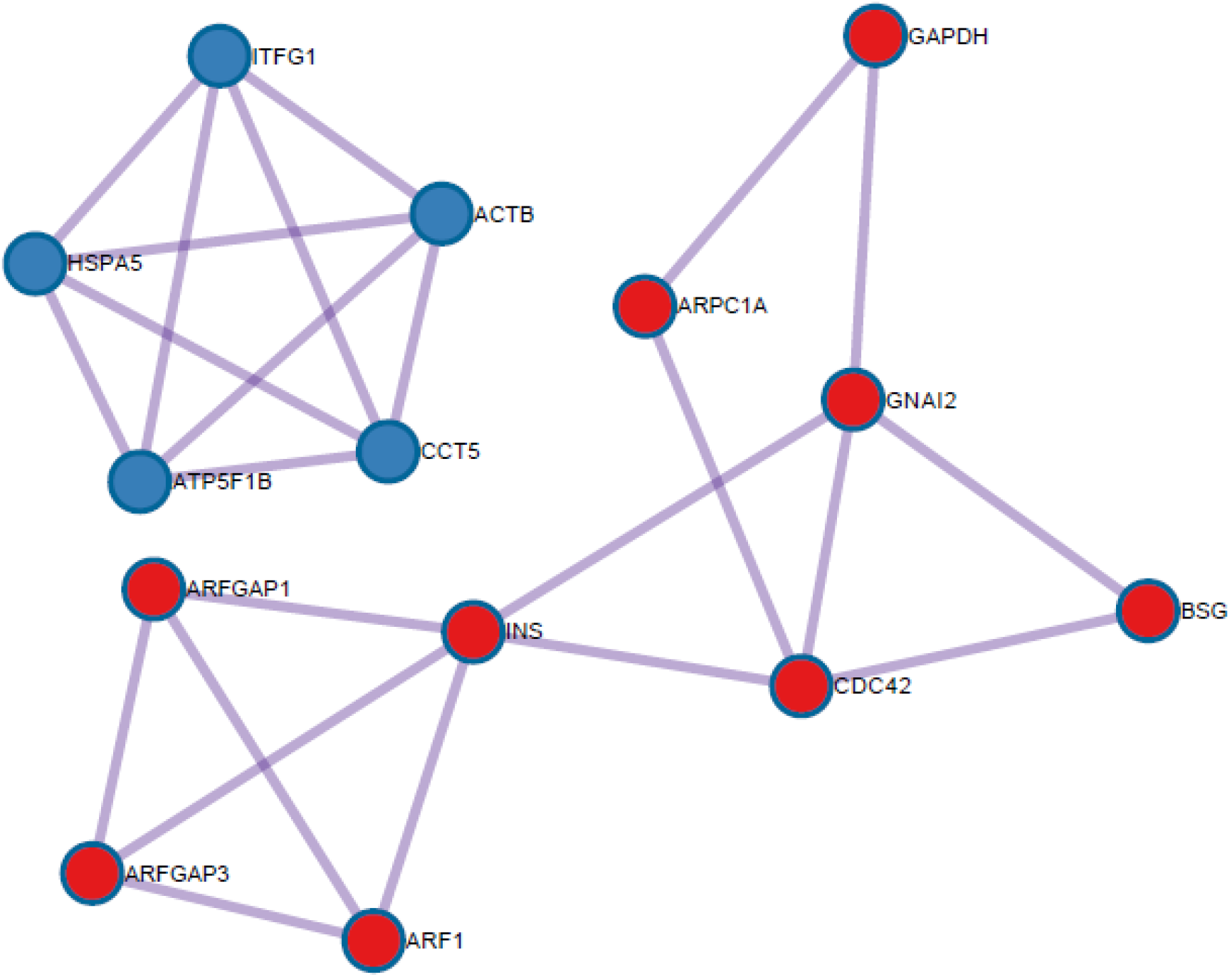
Two highly interconnected regions (clusters) in a protein-protein interaction enrichment analysis according to 95 genes of scsvm2, submitted to metascape.

#### 3.2.3 Dataset3

Table 4 reports the top retrieved terms for each method by Enrichr. It is clear that scsvm2 has more expressed genes (and good ranking results) in pancreatic islet and alpha cell terms when compared to all other methods. In particular, 27 out of 2316 genes were expressed in pancreatic islets. Totally, 20 and 21 genes were expressed in beta and alpha cells, respectively. Seventeen genes (RBPJL, CELA3A, EGR1, PRSS1, CPA1, CELA3B, JUN, SPINK1, CTRB2, CTRB1, GP2, GCG, MTRNR2L8, DLK1, CLPS, ID4, and FOSB) were commonly expressed in beta and alpha cells. In terms of beta cell, sam had 22 expressed genes (KLF10, CELA3A, PRSS1, CDKN1A, SPINK1, KIRREL2, DTNA, KLK1, REG1A, CTRB1, SLC16A12, CEL, SLC39A14, TMC4, SERPINA4, DLK1, NR4A1, SCG5, GNAS, FAT1, CELP, and KCTD16). Lasso ranking was the best in terms of beta and alpha cells; however, it was the worst among all other methods in terms of the number of expressed genes. Although beta cells were ranked 6th when SAM was used, pancreatic islets were ranked 25th, far lower than ranks obtained by scsvm1 and scsvm2, both of which ranked pancreatic islets 1st. Therefore, the SAM ranking, collectively (i.e., by considering the rankings for all items), was worse than those of scsvm2 and scsvm1. These results demonstrate the superiority of scsvm2 in identifying pancreatic cell types within the pancreatic islet.

**Table 4:** Enriched terms in “ARCHS4 Tissues” using Enrichr based on genes obtained from Dataset3, consisting of acinar and delta pancreatic cells. Rank column shows the order of terms when retrieved. Overlap displays the number of common genes between the uploaded gene and those in the term divided by the number of genes in that term. Best results of a method is in bold.

| Method | Rank | Term | Overlap | P-value | Adjusted P-value |
| --- | --- | --- | --- | --- | --- |
| scsvm1 | 1 | PANCREATIC ISLET | 25/2316 | $5.74 \times 10^{-5}$ | $5.75 \times 10^{-3}$ |
| | 10 | BETA CELL | 19/2316 | $1.19 \times 10^{-2}$ | $1.17 \times 10^{-1}$ |
| | 8 | ALPHA CELL | 20/2316 | $5.61 \times 10^{-3}$ | $6.73 \times 10^{-2}$ |
| scsvm2 | 1 | PANCREATIC ISLET | **27**/2316 | $6.36 \times 10^{-6}$ | $6.87 \times 10^{-4}$ |
| | 8 | BETA CELL | 20/2316 | $5.61 \times 10^{-3}$ | $6.73 \times 10^{-2}$ |
| | 6 | ALPHA CELL | **21**/2316 | $2.50 \times 10^{-3}$ | $3.86 \times 10^{-2}$ |
| baseline | 1 | PANCREATIC ISLET | 24/2316 | $1.60 \times 10^{-4}$ | $8.62 \times 10^{-3}$ |
| | 21 | BETA CELL | 15/2316 | $1.32 \times 10^{-1}$ | $5.70 \times 10^{-1}$ |
| | 11 | ALPHA CELL | 18/2316 | $2.37 \times 10^{-2}$ | $2.13 \times 10^{-1}$ |
| limma | 18 | PANCREATIC ISLET | 10/2316 | $6.74 \times 10^{-1}$ | $9.99 \times 10^{-1}$ |
| | 24 | BETA CELL | 9/2316 | $7.85 \times 10^{-1}$ | $9.99 \times 10^{-1}$ |
| | 72 | ALPHA CELL | 6/2316 | $9.70 \times 10^{-1}$ | $9.99 \times 10^{-1}$ |
| sam | 25 | PANCREATIC ISLET | 19/2316 | $1.19 \times 10^{-2}$ | $4.93 \times 10^{-2}$ |
| | 6 | BETA CELL | **22**/2316 | $1.06 \times 10^{-3}$ | $1.90 \times 10^{-2}$ |
| | 12 | ALPHA CELL | 20/2316 | $5.61 \times 10^{-3}$ | $3.36 \times 10^{-2}$ |
| *t*-test | 34 | PANCREATIC ISLET | 13/2316 | $3.04 \times 10^{-1}$ | $8.42 \times 10^{-1}$ |
| | 77 | BETA CELL | 8/2316 | $8.73 \times 10^{-1}$ | $1.00 \times 10^0$ |
| | 76 | ALPHA CELL | 8/2316 | $8.73 \times 10^{-1}$ | $1.00 \times 10^0$ |
| lasso | 3 | PANCREATIC ISLET | 3/2316 | $1.55 \times 10^{-3}$ | $1.44 \times 10^{-2}$ |
| | 2 | BETA CELL | 3/2316 | $1.55 \times 10^{-3}$ | $1.44 \times 10^{-2}$ |
| | 1 | ALPHA CELL | 3/2316 | $1.55 \times 10^{-3}$ | $1.44 \times 10^{-2}$ |

Because scsvm2 achieved good performance results, we uploaded the 95 genes obtained from scsvm2 to Metascape. Figure S2.5 in Supplementary Additional File 2 shows the PPI network obtained by Metascape. In Figure 4, four highly connected subgraphs were extracted from the PPI network in Figure S2.5. These four subgraphs include the following 18 genes: COL1A2, COL1A1, SPARC, COL3A1, SEC24B, CD59, SERPINA1, TMED10, LAMB2, SPP1, HSP90B1, RPS19, EEF1A1, RPL3, RPLP2, RPS12, UBC, and RPS27. Among these genes, Lin et al. [41] suggested that COL1A1 can act as a biomarker and therapeutic target for type 2 diabetes. RPS12 and RPS27 have played a key role in the differentiation and proliferation of pancreatic cancer cells [42].

**Figure 4:**
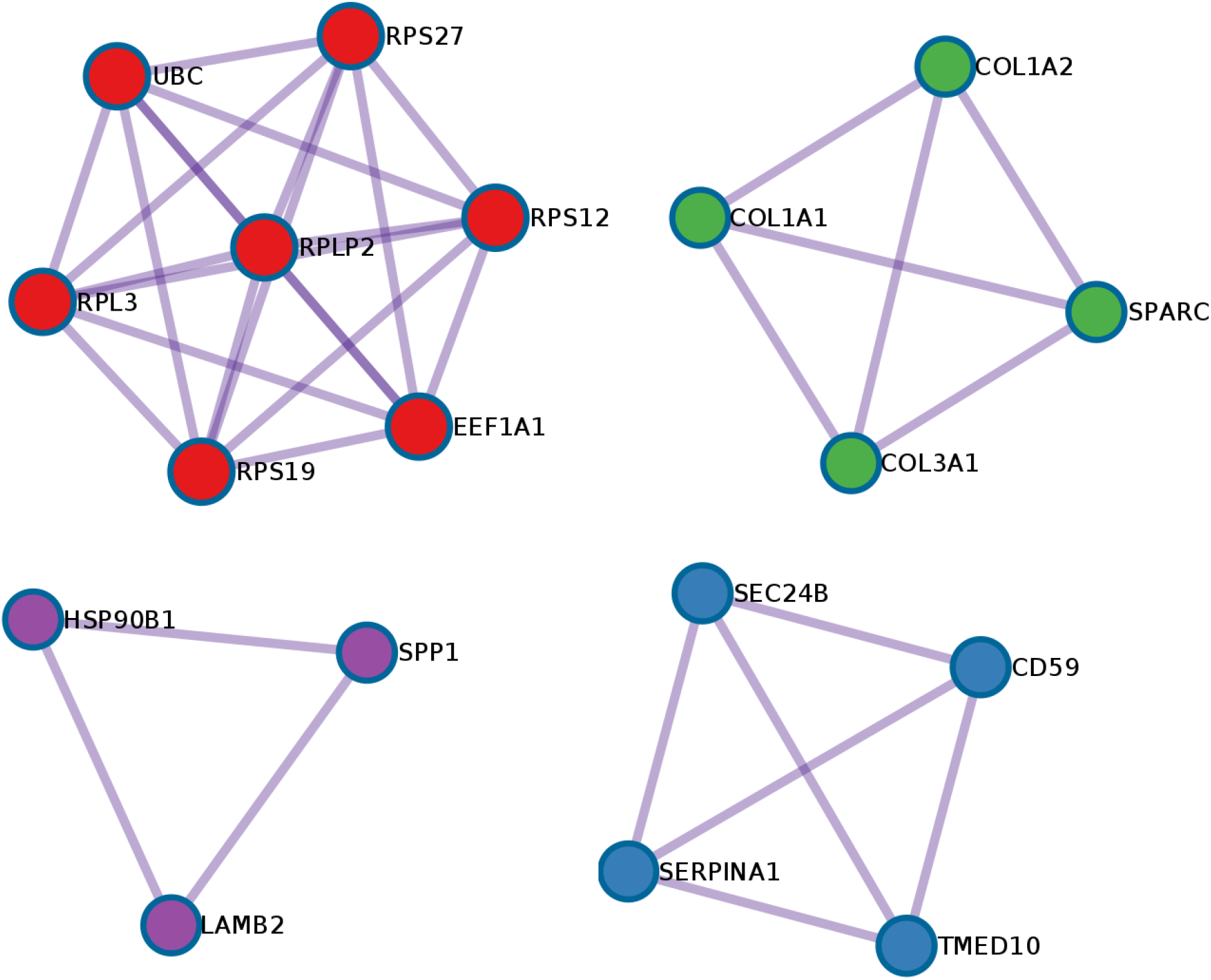
Four highly interconnected regions (clusters) in a protein-protein interaction enrichment analysis according to 95 genes of scsvm2, submitted to metascape.

In Figure S2.6 in Supplementary Additional File 2, we identified seven TFs, including HDAC1, STAT6, RELA, SP3, NFKB1, SP1, and ESR1. Two of these seven TFs were reported in Figure S2.2, namely HDAC1 and SP1. Furthermore, SP3 and NFKB1 have been reported strongly correlated with RAP1 expression, which plays a key role in multiple functions pertaining to pancreatic islet cells [43–45]. For Dataset 3, a list of genes obtained by each method is provided in Supplementary Data Sheet 3. Moreover, a full list of enrichment analysis results of each method is deposited in Supplementary Table 3.

#### 3.2.4 Demonstrating the Discriminative Power

Consider that we are given a dataset pertaining to two cell types (i.e., alpha and beta cells) as shown in Figure 5 (a). Our task is to provide this dataset to a machine-learning algorithm to induce a model, which is then utilized for predicting the types of unseen cells. Figure 5 (c, d) shows that scsvm1 and scsvm2 outperform the baseline (Figure 5 (b)) by learning models that make no errors. Furthermore, scsvm1 (*p-* value = 2.17×10^−3^) and scsvm2 (*p-*value = 5.50×10^−3^) generate statistically significant results (Figure 5 f, g). Because the baseline in Figure 5 (b) has one correctly predicted alpha cell and another incorrectly classified beta cell, no *p*-value was reported, as there were not enough examples in the category. In terms of measuring the classification performance, using the area under curve (AUC) performance measure, it can be shown from Figure 5 (h) that scsvm1 and scsvm2 achieved the highest results, 100%, when compared to the baseline, which generated 75%. These results demonstrate the superiority of the proposed learning algorithms, scsvm1 and scsvm2.

Additionally, assume that we have another dataset as shown in Figure 6, where the task is to provide this dataset to a machine learning algorithm, to generate a model, which then takes new examples to predict the cell type. It can be seen from Figure 6 (d) that the induced model scsvm2 classifies all examples correctly. The second-best model is scsvm1, which incorrectly classifies one out of five beta cells as an alpha cell. In contrast, the baseline was the worst among the three, and incorrectly classified two out of five beta cells as alpha cells. Results obtained by scsvm1 and scsvm2 were statistically significant with *p*-values of 2.846×10^−3^ and 2.842×10^−3^ respectively (Figures 6 (f, g)). Although the baseline generated highly statistically significant results (*p*-value 1.865×10^−5^), the performance result was the worst as it achieved an AUC of 80% which was quite low compared to scsvm1 and scsvm2 which generated an AUC of 90 and 100%, respectively (Figure 6 (h)).

**Figure 6:**
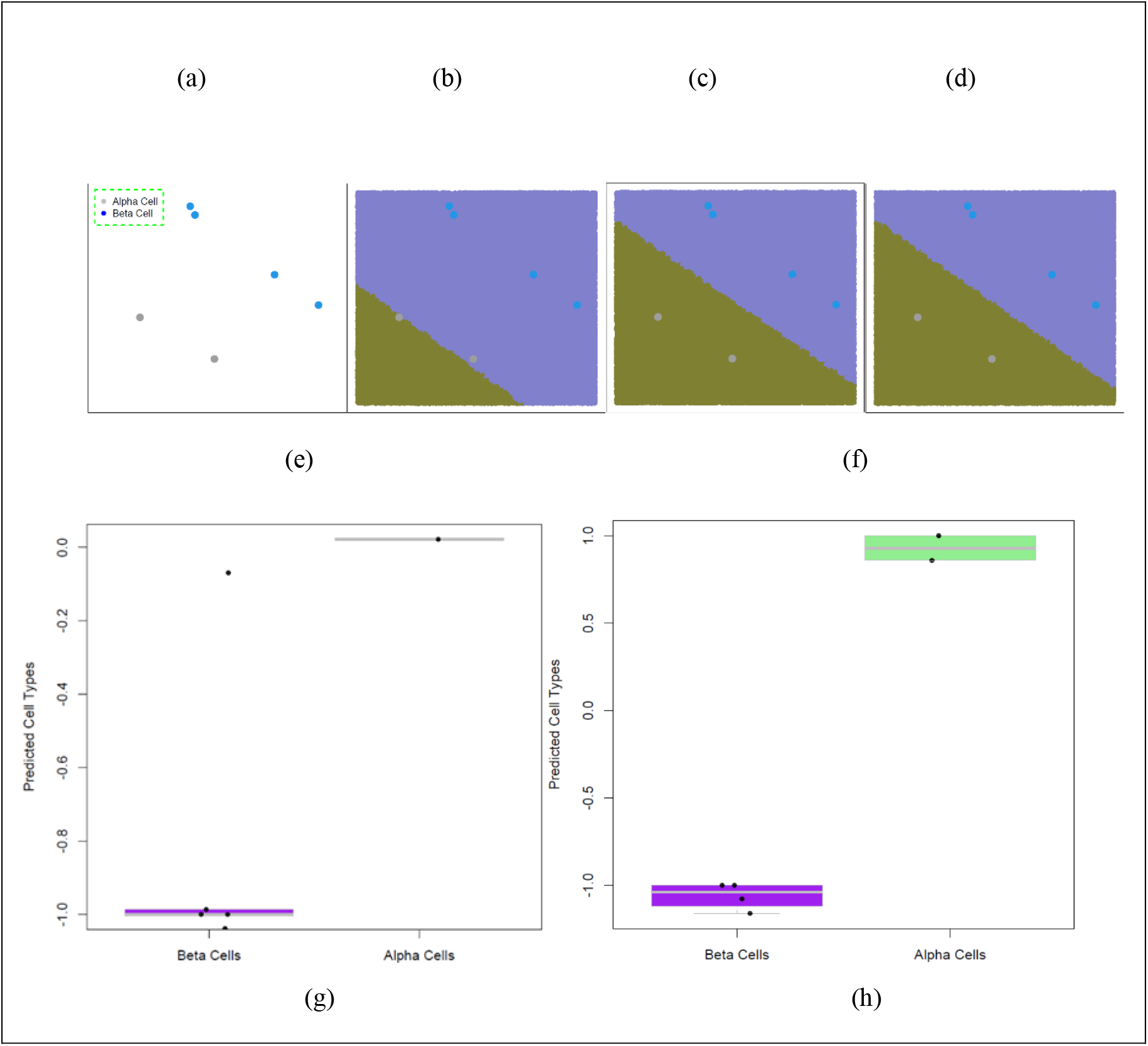
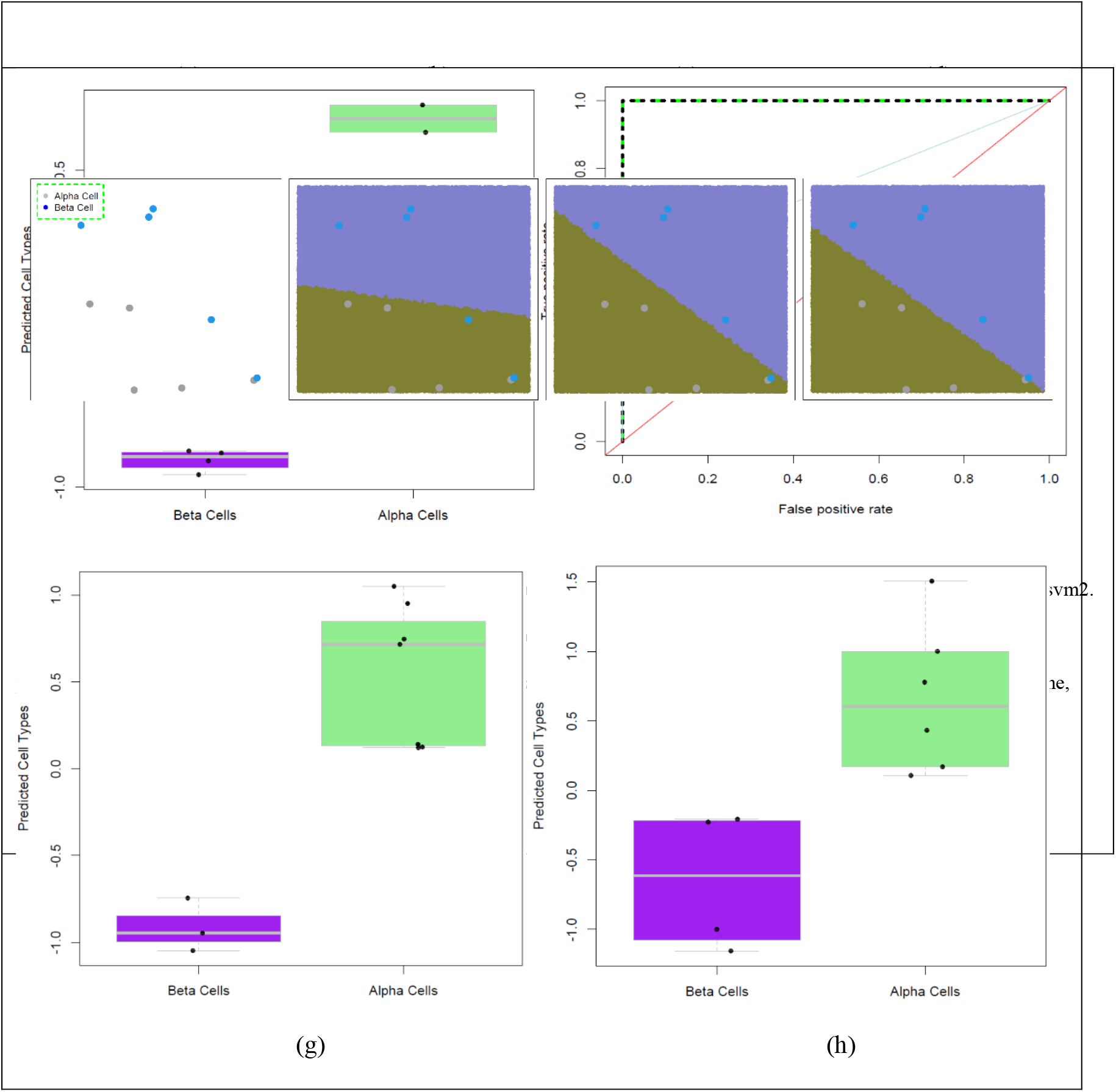
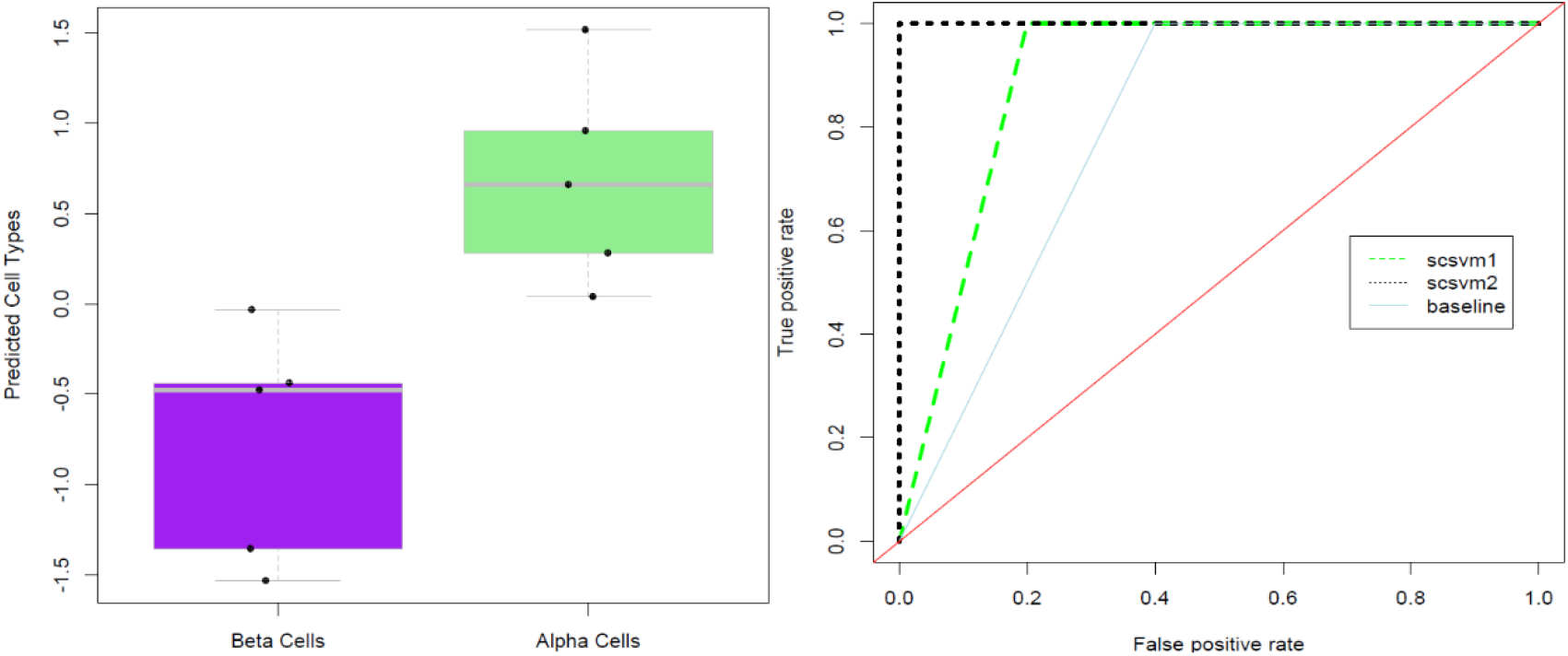
(a) Dataset, (b) decision boundary obtained by the baseline, (c,d) decision boundaries obtained by the scsvm1 and scsvm2, respectively. Prediction results pertaining to alpha and beta cell types using dataset in a. Boxplot and strip charts (e-g) show the differences predicted cell types, beta or alpha, for the baseline, scsvm1, and scsvm2, respectively. (h) shows the ROC curves for the baseline, scsvm1 and scsvm2 by displaying the true positive rate against the false positive rate of each model.

## 4 Discussion

Therapeutic targets pertaining to the pancreas provide a deep biological insight into its complex molecular mechanisms. Therefore, advanced computational methods can aid in speeding up the drug discovery process by providing a basis for the development of drugs for pancreatic disorders. Therefore, we proposed a computational framework to identify influential genes from three expression-profiling datasets. We introduced a new constrained optimization version of SVM and solved an optimization problem using the CVXR package in R, to get weights w, which correspond to the importance of genes in the dataset. The higher the weight, the more important the gene. Then, we selected the top 95 genes and performed enrichment analysis. In terms of cell types, the top retrieved terms by our methods (scsvm1 and scsvm2) consisted of more expressed genes in the pancreatic islet cells, including beta and alpha islet cells. Moreover, from the highly connected subgraphs within the PPI networks and TFs, we identified important genes that have been reported to be related to pancreatic islet functions, diseases, and treatments.

In terms of the results (See Tables 2–4), our computational framework (when coupled with enrichment analysis) acts as a search engine which retrieves the most relevant terms (e.g., pancreatic islet cells) out of many. Much like a search engine, genes and terms can be thought of as contents and pages, respectively. The retrieved terms are related to the selected genes. If the genes in question are not related to pancreatic islet cells, then the retrieved terms are not related to the pancreas. It can be seen from the results that the lower the ranking with more expressed genes, the better the performance. Therefore, the results obtained by our computational framework are better than those obtained from other methods.

Enrichment analysis tools act as search engines for querying gene sets to (1) extract information pertaining to various biological contexts and (2) help bioinformaticians (or experimental biologists) understand biological phenomena [46]. In this study, genes were obtained by a computational method. To understand the functions of candidate genes within the pancreatic context, we used them as inputs for enrichment analysis tools to rank related terms (e.g., pancreatic-related terms). For validation at the protein level, we used Metascape to identify therapeutic targets within the candidate genes and related transcription factors that contribute to pancreatic functions.

For the constrained optimization problem in Equation 1, we set *u* = 0.5, *r* = 0.1, and *c* = 1. The only difference is *k*, which is set to 1 in the scsvm1, while it is set to 2 in scsvm2. v in the function *Φ* is related to 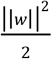 in Equation 1, leading to different objective functions (refer to Equations 2 and 3). In terms of solving the problem, we defined the objective function and constraints, then used the ‘Problem’ and ‘Solve’ functions in the CVXR package, to define and solve the optimization problem [26]. For comparisons among methods, we selected the top 95 genes obtained by each method. The lists of selected genes in Dataset1, Dataset2, and Dataset3 are provided in Supplementary Data Sheet 1, Data Sheet 2, and Data Sheet 3, respectively. There was no significant difference in performance between analyses of 95 and 100 genes. Using the full gene list as an input to Enrichr would be comparable to a long user query in a Google search. However, a list of 95 genes in this study was an acceptable user query length for the ranking of terms using Enrichr. In Dataset 2, It is worth pointing out that SAM and *t*-test provide five and three genes, respectively, for further enrichment analysis. This is because the selection of genes in these methods (including limma) is based on *p*-values < 0.01 (See list of genes in Supplementary Data Sheet 2). Therefore, fewer genes were selected, which would meet this requirement [19, 32]. As lasso shrunk coefficients for genes with weak discrimination power to zero, it provided eight, fifteen, and three genes for Dataset 1, Dataset 2, and Dataset 3, respectively (see list of genes in Supplementary Data Sheet 1, Data Sheet 2, and Data Sheet 3) [47]. Finally, based on the enrichment analysis, our methods performed the best for the retrieved pancreatic cell types in terms of both the ranking and number of expressed genes. The strength of the proposed computational framework relies heavily on the new variant of the constrained optimization problem for SVM. As can be seen in Figures 5-6, our methods generate better performance results when compared to the baseline. In addition, our method is more robust as it finds the maximum marginal hyperplane and conceals the effects of slack variables, which allow for some training errors in the baseline (See Figures 5 and 6). If slack variables are not introduced, hard margins are created in SVM, producing no solution when training data is not linearly separable. Moreover, for Datasets 1-3, the probabilities of any method coinciding with other methods in terms of the 95 selected genes are 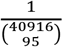, 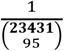, and 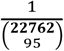, respectively, where 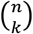 is the binomial coefficient, and *n* and *k* are the number of genes and selected genes, respectively. Therefore, the reproduction of the exact same results as reported in our study is highly unlikely.

One of our aims was to explore therapeutic targets for pancreatic-related disorders. Therefore, we randomly selected three expression profiles pertaining to the pancreas. All of the expression profiles in the datasets were measured by recently developed single-cell technologies. These datasets are related to various pancreatic cell types (as described in Section 2.1) and have been used to understand pancreatic functions and diseases, such as diabetes [23, 24, 48].

In this study, labeled data consisting of expression profiles along with their corresponding pancreatic cell types were used. Column names correspond to genes of interest. We evaluate the labeled data by an SVM algorithm to determine w and b. As SVM prediction is a function of the sign of weighted features (i.e., genes) and bias (i.e., sign(w.x+*b*)), genes with higher weights are important. Our results demonstrate the successful application of SVM to expression profiles generated by single-cell technologies. For deep learning (DL), including neural networks, a similar interpretation to that of SVM is not possible. This is because most DL models do not rely on domain expertise for feature engineering, thereby addressing representation as well as learning. Without domain knowledge, it would be challenging to reach the conclusions obtained in the present study.

Although all of the approaches employed in this study depend on labeled data provided by domain experts, our methods require additional running time to solve the optimization problem to learn w and *b*. When compared to deep learning models, the proposed models provide a simple explanation for selecting important genes in terms of the weight vector w, where higher weights indicate the importance of genes. However, inferences similar to those obtained by SVM may not be feasible for DL models, which are not designed to rely on domain knowledge for feature engineering. This demonstrates an emerging challenge for DL models within the omics domain.

## 5 Conclusion and Future Work

We present a computational framework for identifying influential genes related to pancreatic islet functions and diseases. The framework works by processing expression profiles pertaining to pancreatic islet cells, which are then provided to a new constrained optimization of SVM, to quantify the importance of genes in these transcriptomic expression profiles. Three transcriptome expression profile datasets (GSE87375, GSE79457, and GSE110154) were downloaded from the GEO database. Then, an enrichment analysis was performed on the top *p* selected genes, comparing proposed methods against existing methods, which include the baseline. Results indicate that our methods outperformed all other methods by retrieving the top terms with more expressed genes in pancreatic islet cells. Furthermore, we identified 44 genes and 10 TFs (SP1, HDAC1, EGR1, E2F1, AR, STAT6, RELA, SP3, NFKB1, and ESR1) which have been related to pancreatic functions, diseases and therapeutic targets. The 2D example summarized in Section 3 demonstrates the differences in decision boundaries obtained by our methods and the baseline during the training phase. The relative loss incurred by the decision boundaries for our method was less than that of the baseline; accordingly, our methods outperformed the baseline, as the selection of genes depends on the weights (w) of features in SVM. Notably, prediction by SVM is calculated by the sign of weighted features plus the bias (i.e., sign(w.x+*b*)). Moreover, the proposed methods utilizing the new variant also achieved significant results when compared to the baseline.

We suggest that similar strategies should be implemented to (1) utilize the computational framework to unveil molecular mechanisms and find therapeutic targets in various cells, including the liver, heart, and lung; (2) collaborate with hospitals to improve the poor therapeutic outcomes for diseases; and (3) derive gene signatures from expression-profiling data using our new SVM variant, for assessment in various biological tasks.

## CRediT authorship contribution statement

**Turki Turki:** Conceptualization, Formal analysis, Methodology, Data curation, Software, Supervision, Writing - original draft, Visualization, Investigation. **Y-h. Taguchi:** Conceptualization, Formal analysis, Methodology, Data curation, Validation, Writing - original draft.

## Conflict of interest

None declared.

## Funding

This project was funded by the Deanship of Scientific Research (DSR) at King Abdulaziz University, Jeddah, under grant no. G: 028-611-1442. The authors, therefore, acknowledge with thanks DSR for technical and financial support.

**Turki Turki** received a B.S. in computer science from King Abdulaziz University, an M.S. in computer science from NYU.POLY, and a Ph.D. in computer science from the New Jersey Institute of Technology. He is currently an associate professor with the Department of Computer Science, King Abdulaziz University, Saudi Arabia. His research interests include machine learning, data science and bioinformatics. His research has been published in journals such as *Expert Systems with Applications, IEEE Journal on Selected Topics in Signal Processing*, *BMC Medical Genomics and Computers in Biology and Medicine*. He is an editorial board member of *Sustainable Computing: Informatics and Systems, BMC Medical Genomics, PLOS ONE*, and *Computers in Biology and Medicine*.

**Y-h. Taguchi** received a B.S. degree in physics from the Tokyo Institute of Technology and a Ph.D. degree in physics from the Tokyo Institute of Technology. He is currently a full professor with the Department of Physics, Chuo University, Japan. His works have been published in leading journals such as *Physical Review Letters*, *Bioinformatics*, and *Scientific Reports*. His research interests include bioinformatics, machine learning, and nonlinear physics. He is also an editorial board member of *PloS ONE*, *BMC Medical Genomics*, *Medicine* (Lippincott Williams & Wilkins journal), *BMC Research Notes*, and *IPSJ Transaction on Bioinformatics*.

## Notes

### Competing Interest Statement

The authors have declared no competing interest.

### Summary of Updates

Modification based upon reviewers' comments.

