## Supplementary material for "A new machine learning based computational framework identifies therapeutic targets and unveils influential genes in pancreatic islet cells": Additional File 1.docx

### **S1.1 Support Vector Machines**

An SVM is a machine learning algorithm that employs training data {(x_1_,y_1_),…,( x*_m_*,y*_m_*_)_} comprising *m* labeled examples to induce model *h*. The data is used to perform predictions of unseen examples. In Equation 1, we show the traditional constrained optimization problem for SVM in the non-separable case, where the main goal is to learn arguments (i.e., weights, w, and bias, *b*) that minimize the cost function, subject to linear constraints. This method guarantees that a decision boundary with the maximum margin is obtained. Since training data are not linearly separable most of the time, slack variables ξ*_i_* are introduced to allow for learning w and *b* while incurring loss in the training data. It can be seen from Equation 2 that the model induced by SVM is just the dot product of weights and features in addition to the bias (i.e., w.x + *b*). For a given test example x*, prediction is performed by taking the sign of w.x* + *b*. *I*() as an indicator function that maps to 1 if its arguments are true. If its arguments are false, however, it maps to -1. Therefore, the outcome y^*^ is either +1 or -1, depending on the input class label that maximizes the functional value.

| 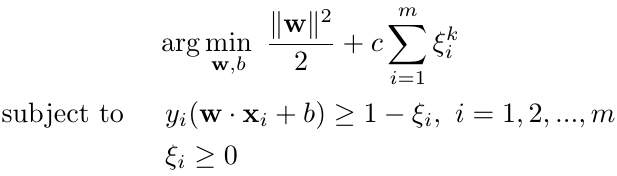 | (S1.1.1) |
| --- | --- |

| 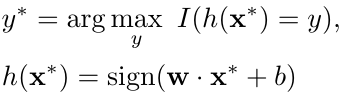 | (S1.1.2) |
| --- | --- |

### **S1.2 Huber Regression (HR)**

HR is a machine learning algorithm that takes input training data {(x_1_,y_1_),…,( x_m_,y_m)_} to induce model *h,* which then takes unseen examples to perform predictions [21]. Unlike the constrained optimization problem of SVM (as seen from Equation S1.1), the optimization problem in HR is unconstrained, aiming to learn weights (also called coefficients) β. Moreover, HR offers different loss functions, as seen from Equation S2.1. For a new test example, prediction is performed by taking the dot product of coefficients and features (i.e., x*. β). The outcome is a continuous value.

| 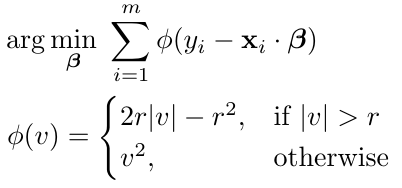 | (S1.2.1) |
| --- | --- |
