## Supplementary material for "A new machine learning based computational framework identifies therapeutic targets and unveils influential genes in pancreatic islet cells": Additional File 2.docx

### Metascape enrichment analysis results for Dataset1

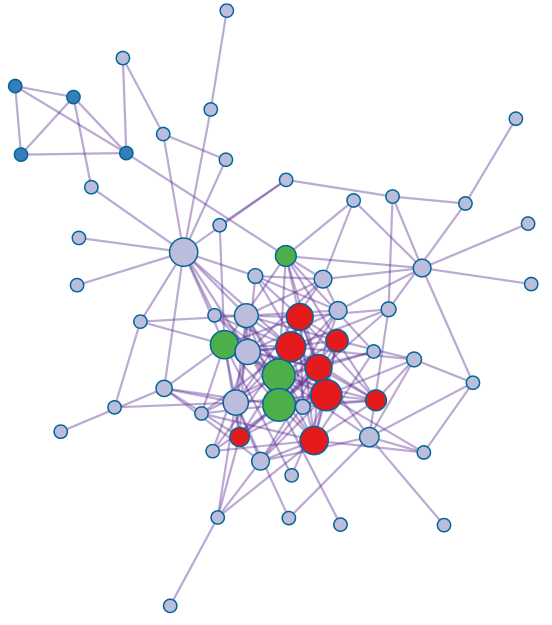

**Figure S2.1**: Protein-Protein interaction network provided by metascape for enriched genes of scsvm1

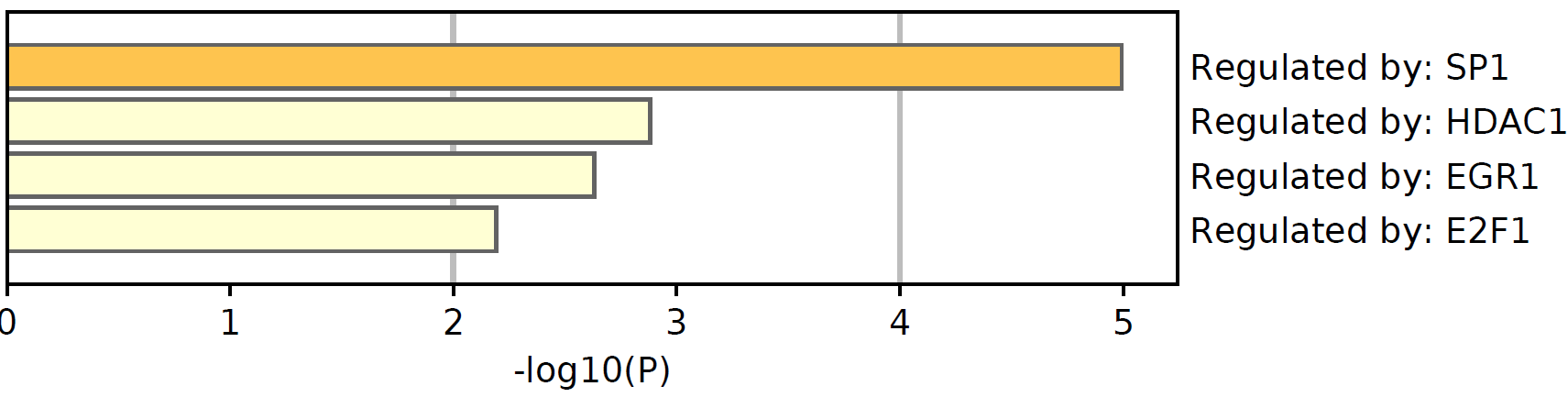

**Figure S2.2**: Four transcription factors regulating the submitted genes by scsvm1 to metascape.

### Metascape enrichment analysis results for Dataset2

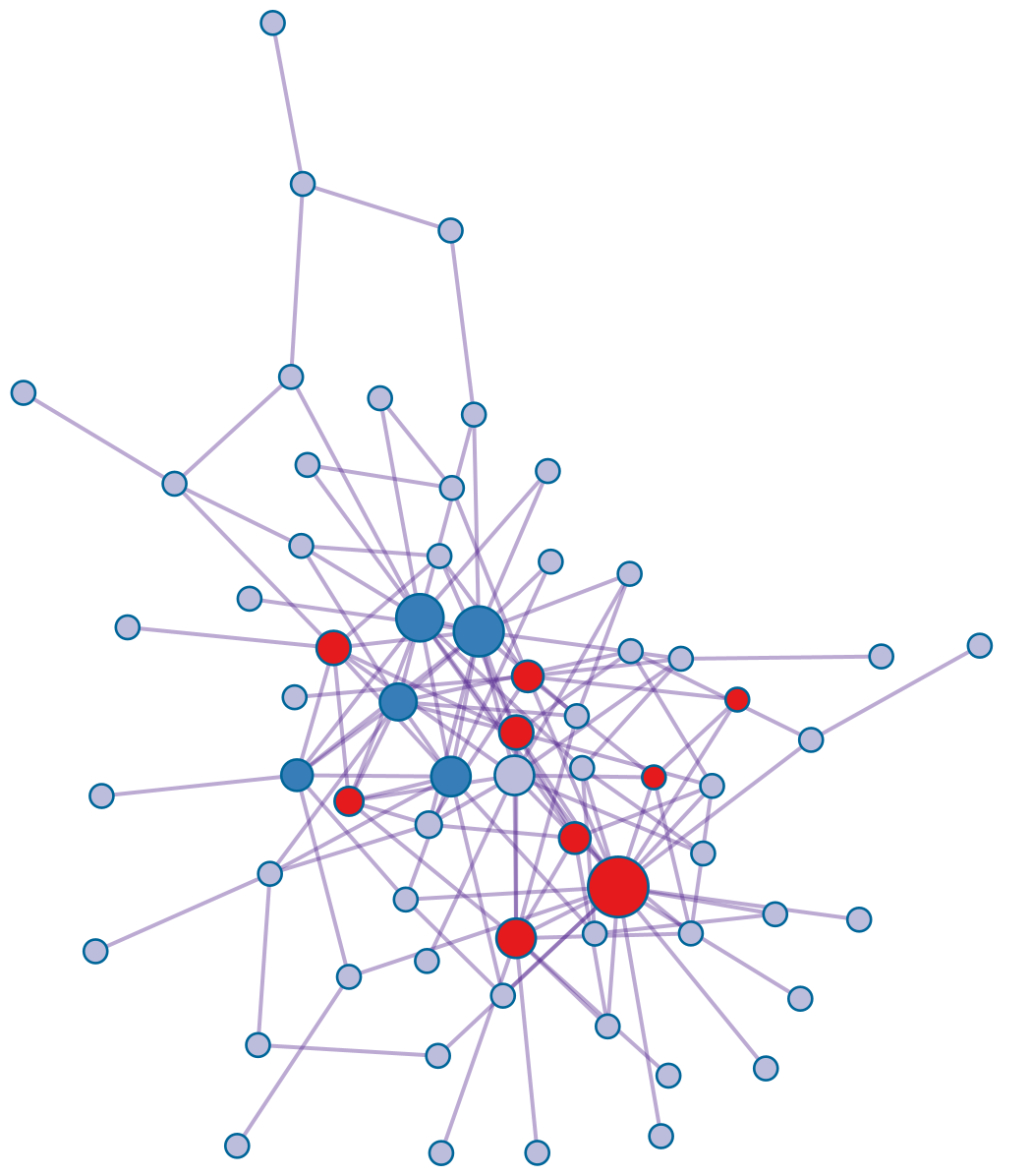

**Figure S2.3**: Protein-Protein interaction network provided by metascape for enriched genes of scsvm2

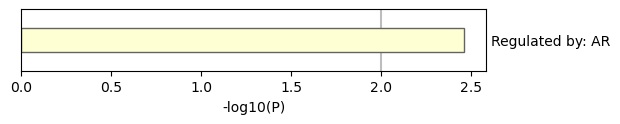

**Figure S2.4**: A Transcription factor regulating the submitted genes by scsvm2 to metascape.

### Metascape enrichment analysis results for Dataset3

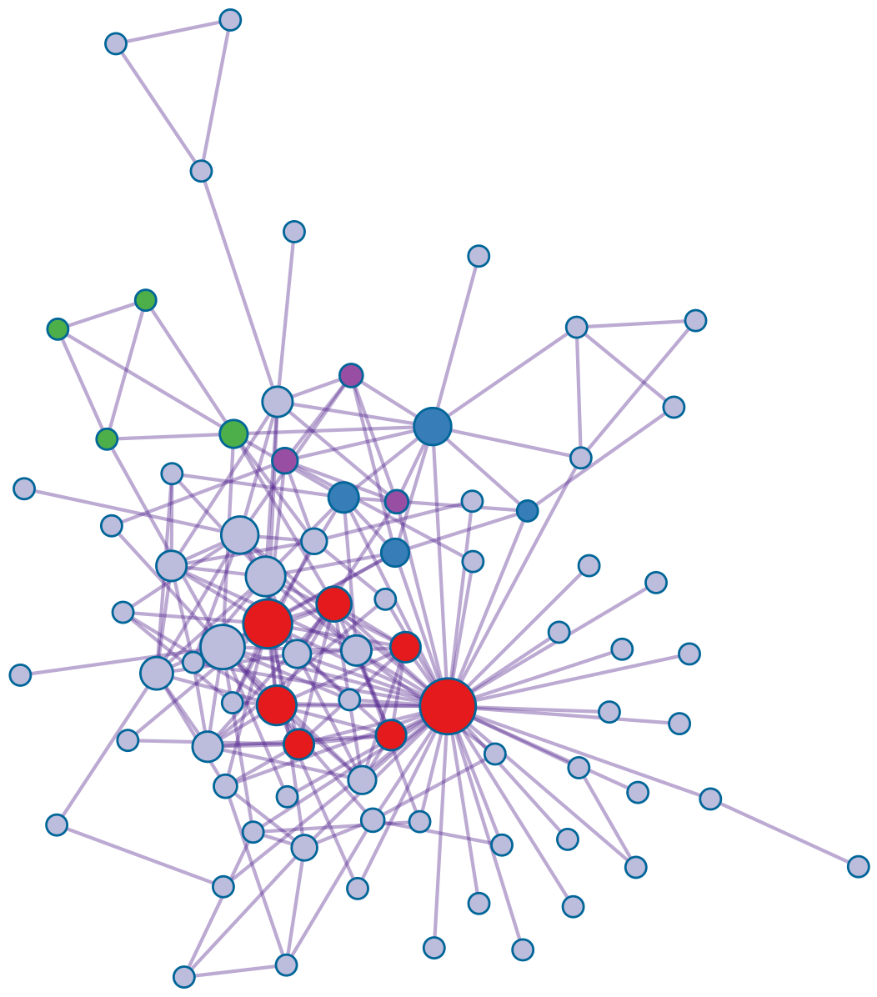

**Figure S2.5**: Protein-Protein interaction network provided by metascape for enriched genes of scsvm2.

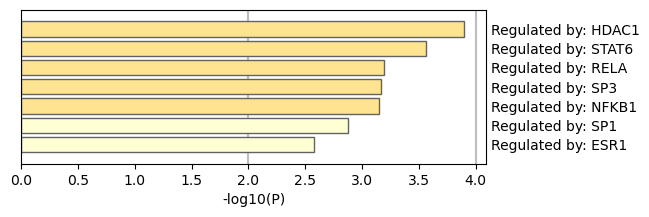

**Figure S2.6**: Seven Transcription factor regulating the submitted genes by scsvm2 to metascape.
